# A soft tick vector of *Babesia* sp. YLG in Yellow-legged gull (*Larus michahellis*) nests

**DOI:** 10.1101/2023.03.24.534071

**Authors:** Claire Bonsergent, Marion Vittecoq, Carole Leray, Maggy Jouglin, Marie Buysse, Karen D. McCoy, Laurence Malandrin

## Abstract

*Babesia* sp. YLG has recently been described in Yellow-legged gull (*Larus michahellis*) chicks and belongs to the Peircei clade in the new classification of Piroplasms. Here, we studied *Babesia* sp. YLG vectorial transmission by ticks in the simplified environment of a single seabird breeding colony where the Yellow-legged gull is the sole vertebrate host, *Ornithodoros maritimus* (syn. *Alectorobius maritimus*) the sole tick species, and *Babesia* sp. YLG is the only blood parasite species detected in chicks of the colony. We collected ticks over four years, maintained certain individuals through moulting or oviposition, and dissected fresh ticks to isolate different organs and test for the presence of the parasite using molecular assays. We report the first strong evidence of a Piroplasmidae transmitted by a soft tick. Indeed, *Babesia* sp. YLG DNA was detected in the salivary glands of nymphs, females and males, a necessary organ to infect for transmission to a new vertebrate host. Parasite DNA was also found in tick ovaries, which could indicate possible transovarial transmission. Our detection of *Babesia* sp. YLG DNA in several male testes and in endospermatophores, and notably in a parasite-free female (uninfected ovaries and salivary glands), raise the interesting possibility of sexual transmission from infected males to uninfected females. Future work in this system will now need to focus on the degree to which the parasite can be maintained locally by ticks and the epidemiological consequences of infection for both *O. maritimus* and its avian host.

## Introduction

The transmission of vector-borne disease is tightly linked to the ecology of both the vertebrate host and the vector species. The circulation of vector-borne infectious agents can be particularly favoured among marine birds due to their tendency to breed in large and dense colonies and the frequent presence of nest-associated ectoparasites (McCoy et al., 2016). Although seabirds are normally only present in the colony during the breeding season, a few months per year, during this time they aggregate by hundreds to thousands, ensuring a short-term but reliable food source for blood-feeding vectors. This is particularly true for ticks that can tolerate a wide variety of abiotic conditions when not associated with a host and can survive long periods without feeding (Dautel & Knulle, 1997; Sonenshine & Roe, 2014). As seabirds tend to show strong fidelity to their breeding colony among years, these colonies can support large tick populations that build up over time (Danchin, 1992).

Seabirds are known hosts for a large variety of nidicolous – or nest-dwelling – tick species. These ticks are well represented by species of the Argasidae family (soft ticks), although some important nidicolous ticks belong to the Ixodidae family (hard ticks) (Dietrich et al., 2011). Between family differences in tick feeding and life histories may profoundly influence pathogen transmission dynamics in these marine systems. Argasid ticks have short and repeated blood meals during nymphal and adult stages, multiple nymphal instars and egg clutches, as well as rapid stage/instar transitions (Vial, 2009). On the contrary, Ixodid blood meals and metamorphoses are usually long and limited to one per life stage (Gray et al., 2014). Therefore, Argasids likely have shorter generation times and more opportunities to acquire and/or transmit pathogens within a seabird breeding season compared to Ixodids, at least at very local scales (Kada et al., 2017).

The pathosystem in the present study includes three component species: a seabird host, the Yellow-legged gull (YLG - *Larus michahellis*, Laridae), parasitized by a soft, nidicolous tick (*Ornithodoros maritimus*, Argasidae, syn. *Alectorobius maritimus*, Mans et al. 2021) and by a blood parasite (*Babesia* sp. YLG, Piroplasmidae). The YLG is a large and widespread species that forms breeding colonies throughout the Mediterranean basin and along the Atlantic coasts of southern Europe and northern Africa (Pons et al., 2004). The breeding season lasts three to four months, between March and July depending on the locality, and nesting areas are reused year after year. As partial migrants, some birds travel widely after breeding, going as far north as the British Iles, whereas others remain in the vicinity of the colony all year round (Souc et al., 2023). These birds tend to remain gregarious even outside breeding, foraging at sea or scavenging for food in dumps or in harbors (Ramos et al., 2009).

The second component in the system, the soft tick *O. maritimus*, infests nesting colonies of various seabird species (e.g., terns, gulls, shags, shearwaters, boobies etc) from southern Great Britain to North Africa (Hoogstraal et al., 1976), although there remains some doubts on the species status for certain host records (Gomez-Diaz et al., 2012; Dupraz et al., 2016). *O. maritimus* is frequently found in YLG colonies of the Mediterranean basin (Dietrich et al., 2011; Dupraz et al., 2016). All life stages of this tick co-occur in the nest (larvae, nymphal instars and adult stages), and feed generally on birds (chicks and adults) at night when the host is quiet. *O. maritimus* is suspected to transmit a great variety of viruses (Meaban, Soldado and West Nile viruses, for example) and bacteria (*Borrelia turicatae, Rickettsia*-like organisms) (Dietrich et al., 2011; Dupraz et al., 2017), although relatively few studies have examined this aspect in detail.

The last component species of the studied pathosystem is *Babesia* sp. YLG (Apicomplexa), a parasite that we characterized previously from blood of YLG chicks (Bonsergent et al., 2022). This species belongs to the Peircei group, a separate and well-defined clade among the Piroplasmidae (Jalovecka et al., 2019; Yabsley et al., 2017), containing only avian infecting piroplasms. We found a high prevalence of the parasite in young chicks and variable levels of parasitemia, with up to 20% infected erythrocytes in some individuals. Based on infection patterns and suspected vectors of other *Babesia*, it is assumed that transmission among hosts occurs in the colony via *O. maritimus*. However, to date, no formal demonstration of this mechanism has been performed.

Here, we examine this question in more detail, focusing on this pathosystem in the particular context of a single YLG colony on the islet of Carteau (Gulf of Fos, Camargue, France). On this islet, there is only one vertebrate host (i.e., a mono-specific breeding colony of YLGs) and only one tick species (*O. maritimus*) (Dupraz et al., 2017; Rataud et al., 2020). In this simplified ecosystem, we analyzed the transmission of *Babesia* sp. YLG by collecting different tick life stages from YLG nests. Ticks were dissected and organs were analyzed separately to detect the presence of piroplasm DNA and to evaluate different transmission pathways.

## Methods

### 1. Study location and tick sampling

Ticks were collected on the islet of Carteau (GPS coordinates: 43° 22′ 39″ N 4°51′ 28″ E) from different nests during four consecutive breeding seasons (2019, 2020, 2021 and 2022), between March and May, and were sent alive to Nantes for analyses. Carteau is a small sandy islet within the Gulf of Fos in southern France with a Mediterranean climate, typically characterized by wet autumns and winters, and warm, dry summers. The local animal ethical committee (APAFIS n°25183-2020090713423689) approved the capture and manipulation of YLG chicks. Permission to access the Carteau colony was provided by Grand Port Maritime de Marseille and the DDTM 13/Service Mer Eau Environnement/Pôle Nature et Territoires (n°13– 2018–02–2-003).

### 2. Tick identification and dissection

The life stage of each *O. maritimus* tick (nymph, adult female, adult male) was determined after collection and all engorged females were isolated in individual collection tubes. In some cases, engorged ticks laid eggs (if female) or moulted into subsequent life stages prior to analyses. Freshly moulted nymphs and females were used to study trans-stadial transmission of the parasites, whereas eggs were used to study transovarial transmission.

All ticks were dissected under a binocular stereo microscope using single-use equipment to avoid contamination. Salivary glands, ovaries, male genitalia, endospermatophores and caeca were recovered and individually frozen at -20 °C in 20 μl PBS 1X until extraction (Fig. 1). One to three successive egg clutches were collected from ovipositing females. Each clutch of 10 to 200 eggs was analyzed as a separate batch and was crushed in a microtube using a sterile pilar before DNA extraction.

**Figure 1.**
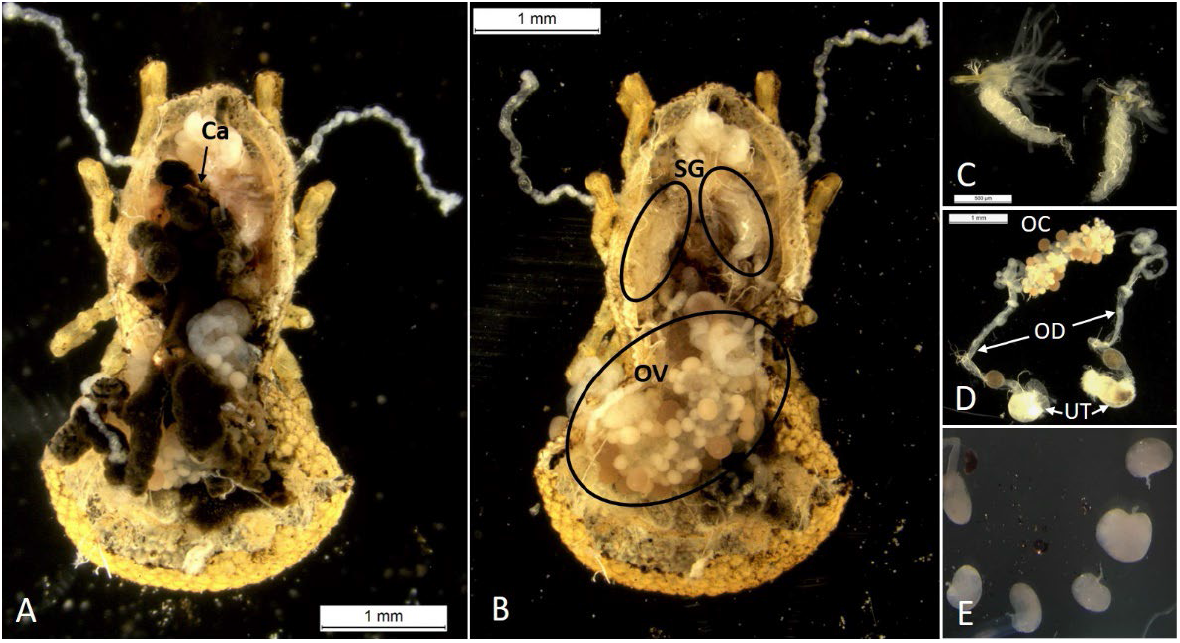
Dissection of an engorged female *Ornithodoros maritimus*. A: all organs visible from the dorsal surface, (Ca: caeca with host blood). B: salivary glands (SG) and ovaries (OV) after the caeca were collected. C: salivary glands. D: ovocytes (OC) at different maturation stages (different sizes and colors), oviducts (OD) and uterus (UT) cut in two parts. E: endospermatophores collected from a female tick (© M. Buysse).

### 3. Genomic DNA extraction from tick organs and eggs

Genomic DNA of tick organs and eggs was extracted using the NucleoSpin Tissue kit according to the manufacturer’s instructions (Macherey-Nagel, Germany). DNA was eluted with 50 μL of elution buffer and was stored at -20°C until use.

### 4. Molecular detection and characterization of piroplasms in tick organs

A nested PCR was performed to detect the 18S rDNA gene of piroplasms in different organs of *O. maritimus*. The primary PCR was conducted to amplify the entire 18S rDNA gene with 5 μL of DNA extracted from the different tick tissues (salivary glands, ovaries, male genitalia, endospermatophores, caeca and eggs), using primers CRYPTOF and CRYPTOR (Table 1) (Malandrin et al., 2010). Reactions were carried out in 30 μL reaction mixtures containing 1 X Go Taq buffer, 4 mM MgCl2, 0.2 mM of each dNTP (Eurobio), 1 unit GoTaq G2 Flexi DNA Polymerase (Promega), 0.5 μM of each primer and 10 μL of DNA template. The amplification conditions comprised 5 min at 95°C followed by 40 cycles at 95°C for 30 s, 30 s at 63°C, 1 min at 72°C, and a final extension at 72°C for 5 min. A nested PCR with primers 18SBp_fw and 18SBp_rev (Table 1) was then carried out with 10 μL of 1/40 diluted amplicons in a 30 μL reaction mixture containing the same components as the first PCR. Cycling conditions were the same as the primary reaction except that the annealing temperature was reduced to 61°C. Amplified fragments were purified with ExoSAP-IT reagent following manufacturer’s recommendations (Affymetrix) and were sequenced bi-directionally using the same primers (Eurofins Genomics, Germany). Sequences were then assembled using the Geneious R6 software (https://www.geneious.com) and an online BLAST (National Center for Biotechnology Information) was performed.

**Table 1.**
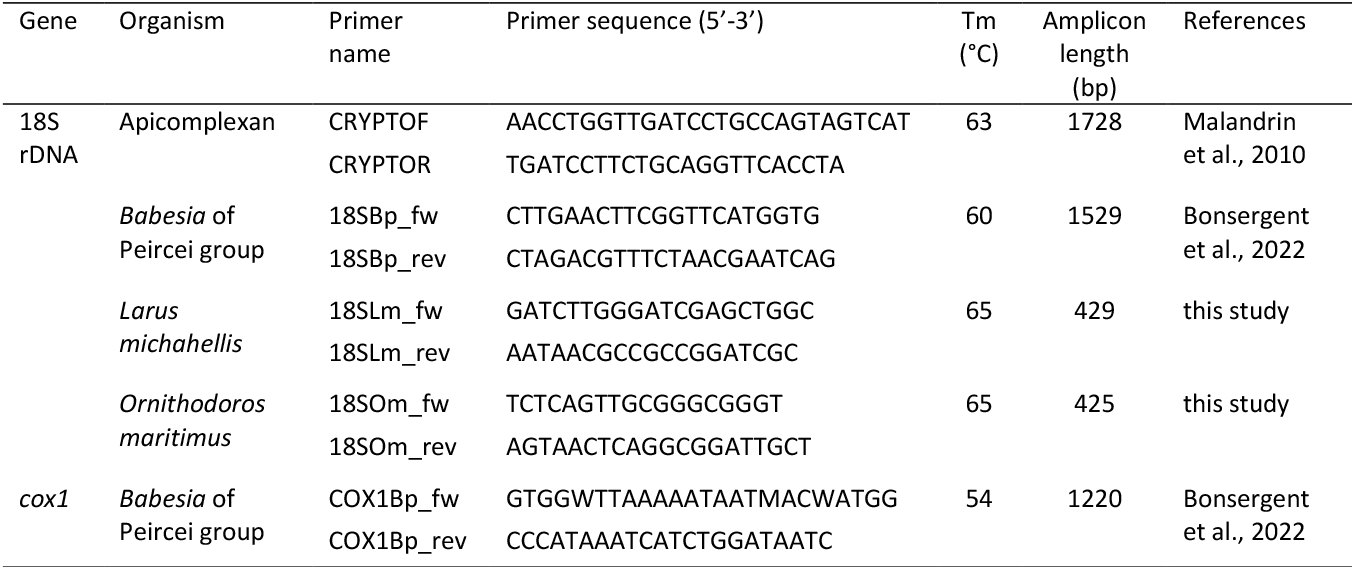
Primers used to amplify 18S rDNA and *cox1* (cytochrome c oxydase subunit 1) genes in the different organisms used in the study.

In order to develop specific primers for controls, the 18S rDNA gene sequences of *L. michahellis* (OP542589, this study) and of *O. maritimus* (OP542591, this study) were aligned with *Babesia* sp. YLG sequence (MZ541058) using Clustal Omega (https://www.ebi.ac.uk/Tools/msa/clustalo/). Regions specific to each organism were selected to design the control primers as described in Supplementary Material 1.

To control for the amount of tick DNA extracted from individual organs, we designed the primer pair 18SOm_fw and 18SOm_rev (Table 1) to amplify a 18S rDNA gene fragment of tick DNA. Reactions were carried out in 30 μL reaction mixtures containing 1 X buffer, 4 mM MgCl2, 0.2 mM of each dNTP (Eurobio), 1 unit GoTaq G2 Flexi DNA Polymerase (Promega), 0.5 μM of each primer and 5 μL of total genomic DNA. PCR cycling comprised 5 min at 95°C, 40 cycles of 30 s at 95°C, 30 s at 65°C, 30 s at 72°C, and a final extension at 72°C for 5 min. This PCR resulted in a 425 bp fragment.

We also attempted to amplify the 18S rDNA gene of the YLG for each *Babesia* sp. YLG positive sample to control for a potential contamination of tick organs by the caecal content that might have occurred during the tick dissection, and consequently the potential presence of *Babesia* sp. YLG DNA coming from the host blood found in the tick gut. As avian red blood cells are nucleated, the contamination of tick organs by caecal spillover should be easily detected. Surprisingly, the CRYPTOF and CRYPTOR primers were able to amplify the 18S rDNA gene sequence of *L. michahellis* (1831 bp) from YLG DNA. Therefore, the control amplification of the 18S rDNA gene of *L. michahellis* was conducted by nested PCR with the use of 18SLm_fw and 18SLm_rev primers (Table 1) after a primary amplification with CRYPTOF and CRYPTOR using the same reaction and cycling conditions as for the 18S rDNA gene of piroplasms, but with an annealing temperature at 65°C and an elongation time of 30 s. This PCR resulted in a 429 bp fragment.

Ticks were considered infected when at least one of their organs was found positive by nested PCR. A tick can be found positive either because it ingested infected blood just before sampling (i.e., parasites in the caeca) or because the parasite had invaded other organs such as salivary glands or ovaries. In the second case, the parasite would potentially be transmissible to a new host during the next bloodmeal or to the next tick generation if trans-ovarial transmission takes place.

### 5. Cox1 gene

To verify that the same genetic variants described in YLG chicks were also found in *O. maritimus*, and to evaluate the possibility of mixed infections in ticks, a fragment of the *cox1* gene was amplified and sequenced from several tick organs. Two successive amplifications of the *cox1* gene of *Babesia* sp. YLG were performed with the primers, COX1Bp_fw and COX1Bp_rev (Table 1, Bonsergent et al., 2022), for *Babesia* positive DNA samples. Both reactions were carried out in 30 μL reaction mixtures containing 1 X buffer, 4 mM MgCl2, 0.2 mM of each dNTP (Eurobio), 1 unit GoTaq G2 Flexi DNA Polymerase (Promega), 0.5 μM of each primer. The template was 5 μL of genomic DNA for the first PCR and 5 μL of 1/20 diluted amplicon for the second PCR. For both amplifications, the cycling conditions comprised 5 min at 95°C, 40 cycles of 30 s at 95°C, 30 s at 54°C, 1 min 20 s at 72°C, and a final extension at 72°C for 5 min. Amplified fragments (1220 bp) were purified, sequenced and analyzed manually.

### 6. Statistical analyses

We used Fisher Exact tests to evaluate variation in infection status among years and life stages (R Core Team, 2023).

## Results

### 1. Tick collections

Over the four field seasons, a total of 144 ticks were collected for analyses: 23 in 2019, 63 in 2020, 38 in 2021 and 20 in 2022. All ticks were found in active nests during standardised searches (see Dupraz et al. 2017). Most were females (107), but males (15) and nymphs (22) were also sampled (Fig. 2). Four females and eight nymphs were freshly moulted. These ticks came from more than 38 different nest sites (6 in 2019, 26 in 2020, 6 in 2021, unknown for 2022), and represent a random sample of ticks from each sampled nest. Among the analyzed ticks, one to twelve came from the same nest. The average number of ticks analyzed per nest was 3.8 in 2019, 2.4 in 2020 and 6 in 2021. In 2022, the nest origin was not recorded.

**Figure 2.**
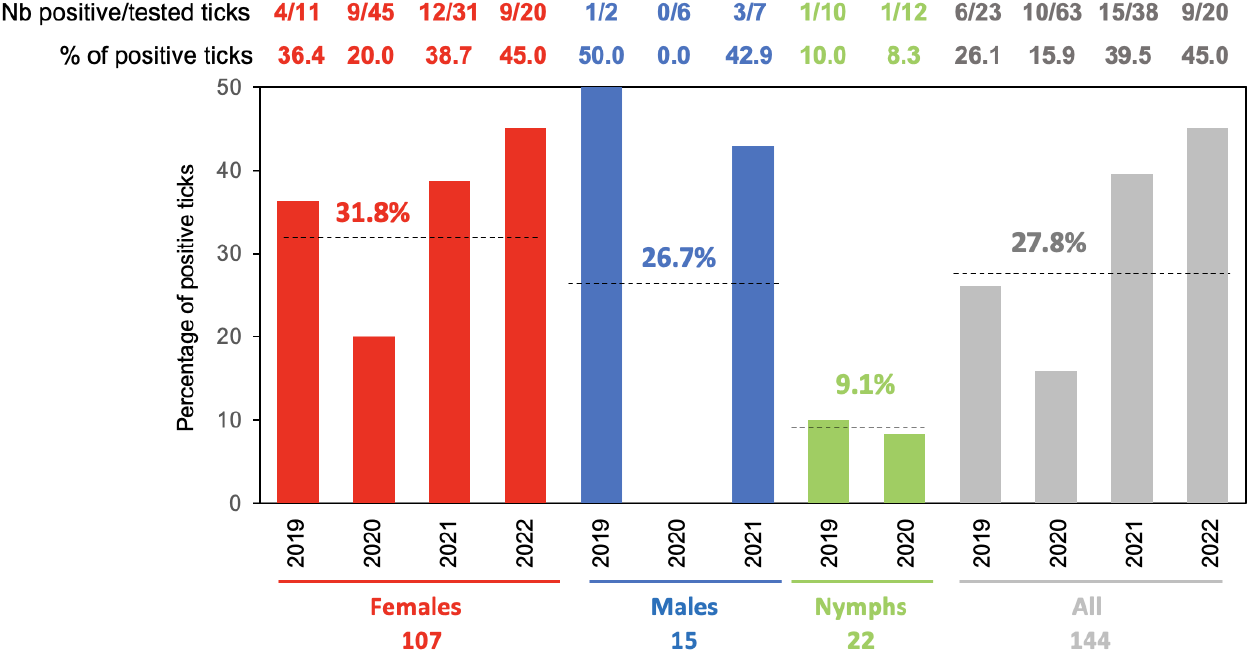
Percentage of *Ornithodoros maritimus* ticks carrying *Babesia* sp. YLG DNA in at least one of their collected organs (salivary glands, ovaries, caeca, endospermatophores from females and/or male genitalia) for each sex/stage and year of collection. The average percentage over a year is indicated by a dashed line with the associated prevalence estimate.

### 2. Prevalence of *Babesia* sp. YLG infected ticks

The 18S rDNA gene of *O. maritimus* was detected in all DNA extracts from tick organs. Of the 144 collected ticks, 40 were found positive for *Babesia* in at least one organ, representing 27.8% of the tested individuals. Infection was present in all years: 26.1% in 2019 (6/23 tested ticks), 15.9% in 2020 (10/63 tested ticks), 39.5% in 2021 (15/38 tested ticks) and 45% in 2022 (9/20 tested ticks) (Fig. 2). Although prevalence varied significantly among years (p-value= 0.01571), this result should be taken with caution because the delay between sampling and testing differed over time (notably due to the covid pandemic) and may have affected detection probability. All tick stages were found infected, with adults tending to show higher prevalence than nymphs (p-value = 0.07875): 31.8% of females (34/107), 26.7% of males (4/15) and 9.1% of nymphs (2/22). Positive ticks came from 18/38 of the identified nests (4/6 in 2019, 9/26 in 2020, 5/6 in 2021), indicating a broad distribution of *Babesia* sp. YLG within the Carteau colony.

### 3. *Babesia* sp. YLG in tick organs and potential transmission routes

#### 3.1. Transmission to the vertebrate host

A total of 137 salivary glands were collected (females-103, males-12, nymphs-22) and analyzed for the presence of *Babesia* sp. YLG (Fig. 3; Supplementary Material 2). In 17 of these (12.4%), the parasite was detected without traces of gut contamination (i.e., the PCR for *L. michahellis* 18S rDNA was negative). Prevalence of positive salivary glands did not vary significantly among life stages (p-value = 0.6006): females (15/103, 14.6%), males (1/12, 8.3%) and nymphs (1/22, 4.5%).

**Figure 3.**
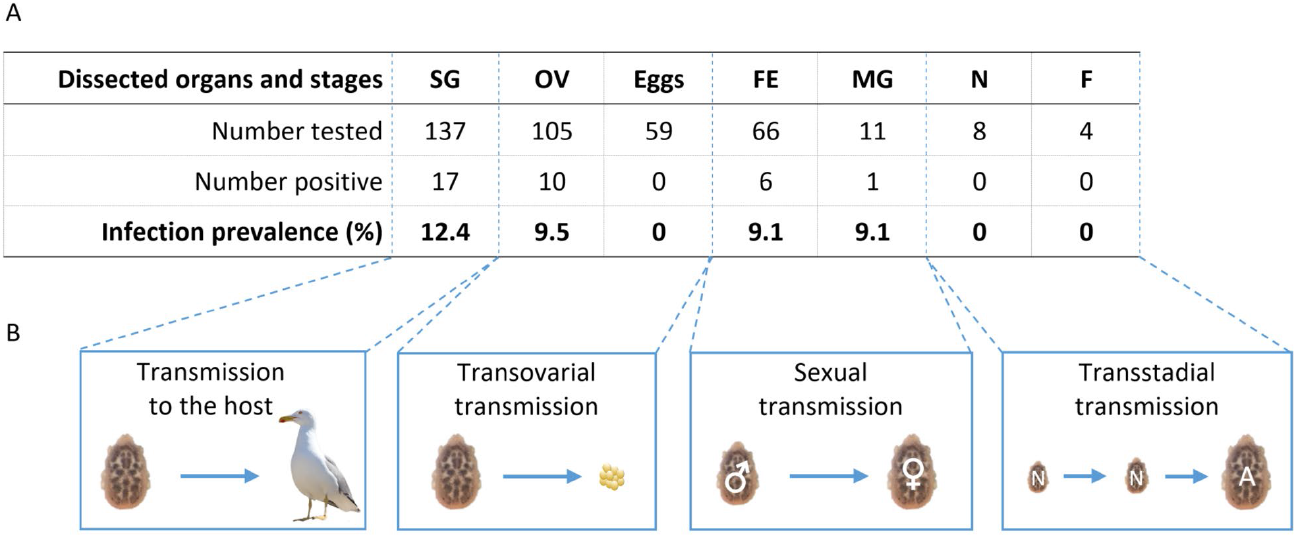
Schematic representation of possible routes of transmission of *Babesia* sp. YLG by *O. maritimus* and the organs analyzed to evaluate each possibility. Panel A: number of tested and positive samples. SG: salivary glands, OV: ovaries, Eggs: egg batches containing 10-200 eggs from a given female, FE: female-derived endospermatophores, MG: male genitalia, N: newly moulted nymphs, F: newly moulted females. Panel B: possible routes of transmission that require the infection of the corresponding tissues in panel A. N: nymphs, A: adult.

#### 3.2. Transovarial transmission

Ovaries from 105 females were recovered and *Babesia* sp. YLG was detected in 10 of them (9.5%), without traces of gut contamination (Fig. 3). A total of 59 egg clutches laid by 46 engorged females were also analyzed. However, no clutch was found positive, even for the three females (and five clutches) for which the ovaries tested positive. Transovarial transmission could therefore not be demonstrated.

#### 3.3. Sexual transmission

Endospermatophores were frequently found in dissected females (66/107 females) indicating recent copulation. *Babesia* sp. YLG was detected in six of them (9.1%) without traces of gut contamination. Positive endospermatophores sometimes came from females (3) for which all other organs tested negative for the presence of *Babesia* sp. YLG (Fig. 3). Among the genitalia tested from 11 males, *Babesia* sp. YLG was detected in only one (9.1%), a prevalence of infection similar to that obtained for the female-derived endospermatophores.

#### 3.4. Transstadial transmission

From the 2020 collection, 12 newly moulted ticks (4 females and 8 nymphs) were tested for the presence of *Babesia* sp. YLG, but the parasite was not detected in any of the dissected organs (Fig. 3).

### 4. Variability of the *cox1* gene

The partial *cox1* gene fragment was successfully amplified from 33 of the 60 organs in which the *Babesia* sp. YLG 18S rDNA gene was detected (17 salivary glands, 10 ovaries, 6 female endospermatophores, 1 male genitalia and 26 caeca), and of these, 28 sequences were obtained from different ticks with lengths comprised between 314 and 1172 bp. Haplotypes a, b, c, d and e, previously described from YLG chicks (Bonsergent et al., 2022), were found either alone (13 ticks) or in mixtures within the same tick (4 ticks), indicating co-infection with different isolates of *Babesia* sp. YLG (Table 2). The same haplotype was often characterized from several organs of the same tick (i.e. ticks 24, 40, M1, M10 and M11). The presence of different combinations of haplotypes were sometimes suggested by double peaks on the chromatograms at a SNP position (i.e. 194bF1 salivary glands and M6 caeca), but we were unable to characterise the co-infecting isolates in these instances (Table 2).

**Table 2.**
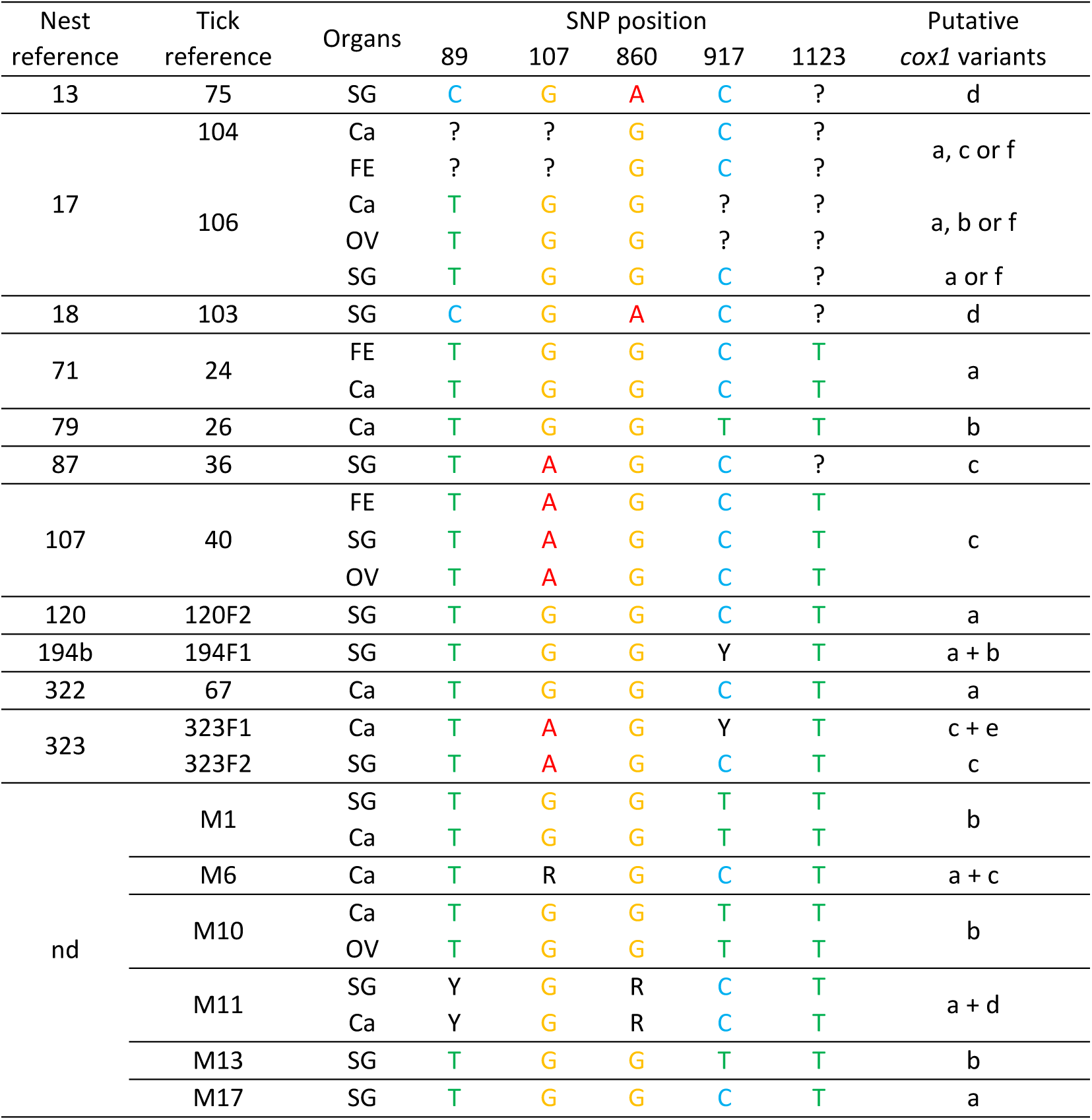
Genetic variants of *Babesia* sp. YLG *cox1* gene found in dissected organs of *Ornithodoros maritimus*. The position of each SNP is indicated as in Bonsergent et al., 2022. FE: female-derived endospermatophores, Ca: caeca, OV: ovaries, SG: salivary glands, ?: undetermined, Y: T or C, R: G or A.

### 5. GenBank deposit

Nucleotide sequences obtained in this study were submitted to Genbank and given accession numbers OP566884 for 18S rDNA and OP588178-OP588182 for *cox1* variants of *Babesia* sp. YLG found in *O. maritimus. L. michahellis* and *O. maritimus* (syn. *Alectorobius maritimus*, Mans et al., 2021) partial 18S rDNA sequences were also deposited under accession numbers OP542589 and OP542591, respectively.

## Discussion

To establish a species as a competent vector for a pathogen, formal proof of acquisition and transmission of the parasite both to and from the host must be demonstrated. However, this task requires experimental infection of pathogen-free individuals under controlled conditions. This type of experiment is particularly challenging for ticks as they are often difficult to maintain and infect under laboratory conditions. In addition, these procedures require the availability of *in vitro* cultures of the parasite and the use of experimental animals, at least as blood donors to cultivate the parasite and feed tick stages. These constraints may prove almost impossible in the case of ticks specialised on wildlife, such as in the biological system we examine here involving a nidicolous soft tick species and a seabird host.

To avoid these issues, we studied *Babesia* sp. YLG transmission in the rather unique and simplified environment of a single seabird breeding colony where the Yellow-legged gull is the sole vertebrate host (so the only source of pathogens for ticks), *O. maritimus* the sole tick species (so the only possible tick vector), and *Babesia* sp. YLG the unique blood parasite species detected in gull chicks (Bonsergent et al., 2022; Dupraz et al., 2017; Rataud et al., 2020). In this environment, we were certain of the active transmission of *Babesia* sp. YLG to chicks. In our previous study, we were indeed able to follow the infection and fulminant multiplication of *Babesia* sp. YLG in two chicks initially found negative or with almost undetectable parasite levels (< 0.1%), and with parasitemia as high as 15-20% recorded two weeks later (Bonsergent et al., 2022). This demonstrated active chick infection in the nest, most probably by *O. maritimus*, an abundant tick species in the colony (Dupraz et al., 2017).

In this pathosystem, we studied *Babesia* sp. YLG transmission by *O. maritimus* by analyzing its presence in dissected tick organs using molecular amplifications of its DNA. This method raises the risk of false positives due to the contamination of organs by infected red blood cells from the host that are present in the tick gut during dissection. However, soft tick blood meals are smaller in volume compared to hard ticks (Vial, 2009), and the risk of spill over during dissection is lower. Nonetheless, to control for this risk and ensure that positive tick organs were due to active tick infections by the parasite and not infected host blood spillover from the gut content, we attempted to detect *Larus michahellis* DNA in all positive tick organ extracts. As avian red blood cells are nucleated, this test represents a very sensitive internal control of gut content contamination. Therefore, in our study, each tick organ that we considered positive (detection of *Babesia* sp. YLG DNA) was negative for *L. michahellis* DNA. Under these conditions, the presence of *Babesia* sp. YLG DNA in salivary glands or ovaries is a strong indication of the passage of the parasite from an initial infection in the gut to other organs that play a role in transmission.

Thus, we report the first strong evidence of the transmission of a Piroplasmidae, namely *Babesia* sp. YLG, by a soft tick, *O. maritimus*. Indeed, *Babesia* sp. YLG DNA was detected in salivary glands, a requirement for transmission to a new vertebrate host. Salivary glands from females, males, and nymphs were positive, indicating a role of these three life stages in the transmission to the host. We were not able to demonstrate transstadial transmission in this study as we could not detect *Babesia* sp. YLG DNA in ticks after moulting. However, we analyzed only 12 moulted ticks of unknown infection status. Future work will need to address this point to better understand transmission efficiency in the system and should also attempt to demonstrate the presence of living parasites in the ticks, rather than working from DNA only.

Current molecular phylogenies now separate Piroplasms into ten clades with major biological differences in life cycles (Jalovecka et al., 2019). *Babesia* sp. YLG belongs to the Peircei group (clade V according to Jalovecka et al., 2019). The life cycles of well-known piroplasmids (*Babesia* sensu stricto or *Theileria* sensu stricto, clade X) have been studied in detail and key biological features such as the vector species, the primary target cells in the vertebrate host, transstadial and transovarial transmission are well described (Jalovecka et al., 2018). These features are poorly known in other clades (II to VI), including the Peircei clade V to which *Babesia* sp. YLG belongs. Phylogenetic work has further suggested that the Percei clade V is separated into two main clusters, one with *Babesia* infecting seabirds and the other with *Babesia* related to terrestrial birds (Grey heron and European roller) (Chavatte et al., 2017; Palomar et al., 2021; Yabsley et al., 2017). One of the *Babesia* described in this second cluster was characterized from a soft tick *Argas* sp. collected in a European roller nest in Spain (Palomar et al., 2021). The only other *Babesia* species reported to be potentially transmitted by a soft tick is *B. vesperuginis* from the Piroplasmidae clade III (Western clade), with *Argas vespertilionis* as the strongly suspected vector (Hornok et al., 2017; Liu et al., 2018; Jalovecka et al., 2019). Up to now, and to our knowledge, transmission of a *Babesia sensu stricto* species (clade X) by a soft tick has never been demonstrated, and could be a biological feature specific to piroplasmids of other clades (III and V). Soft ticks (*Otobius megnini*) collected from naturally infected cattle have been found to carry *Babesia bovis* (clade X), but transmission has not been studied (Malhobo et al., 2021).

Several studies on the characterization of seabird piroplasms from the Peircei clade proposed *Ornithodoros capensis* as a possible vector (Paparini et al., 2014; Peirce & Parsons, 2012; Work & Rameyer, 1997; Yabsley et al., 2009, 2017). *O. capensis* sensu stricto is part of the *O. capensis* complex, like *O. maritimus*, and has been recorded on a range of seabird species breeding close to the equator (Dietrich et al., 2011; Dupraz et al., 2016; Hoogstral et al., 1976). *Babesia*-positive ticks from *O. capensis* females and nymphs were collected from a red-billed gull *Chroicocephalus novaehollandiae* (Paparini et al., 2014) and *O. capensis* larvae were found on a *Babesia*-free chick of *Phalacrocorax capensis* which was found to be infected with *B. ugwidiensis* a week later (Peirce & Parsons, 2012). These indirect observations on phylogenetically closely related tick-pathogen pairs, support a role for *O. maritimus* in the transmission of *Babesia* sp. YLG during the bloodmeal and the possible transovarial transmission of the parasites during tick oogenesis.

Indeed, in addition to finding *Babesia* sp. YLG DNA in the salivary glands, we also detected its presence in tick ovaries, supporting its possible transovarial transmission. However, we were not able to detect *Babesia* sp. YLG DNA in the eggs from infected female ticks. This could be due to the low number of parasites per egg precluding amplification, a probable low number of infected eggs per clutch, and/or to the tick oocyte maturation stage at the time of infection which may or may not allow parasite penetration into the developing eggs (Denardi et al., 2004; Mitchell et al., 2019). Additional analyses will now need to focus on this possible transmission pathway to determine whether *Babesia* sp. YLG can be transmitted vertically in its soft tick vector and thus whether ticks can maintain local infection rates without the required presence of infected host birds. As outlined above, transovarial transmission of piroplasms has only been demonstrated to date in the *Babesia* sensu stricto clade X. This could be a crucial feature in *Babesia* sp. YLG transmission by the soft tick *O. maritimus*. Soft tick females are able to support long periods of starvation, with dormancy behavior, allowing them to wait for the return of breeding birds the following year (Gray et al., 2014). *O. maritimus* females probably feed and oviposit at the onset of spring when adult birds return to their nesting sites. Due to the duration of pre-oviposition and embryogenesis, larvae should appear and be ready for feeding when chicks hatch (about 25 days after egg laying in the case of *Larus michahellis*). If *Babesia* sp. YLG is indeed transovarially transmitted, chicks could be infected very early in their development; larvae are frequently collected feeding on 5 to 39-day-old chicks (Estrada-Pena et al., 1996).

Our detection of *Babesia* sp. YLG DNA in several male testes, along with its presence in endospermatophores, and notably in a parasite-free female (uninfected ovaries and salivary glands), raises the possibility of sexual transmission from infected males to uninfected females. Whether sexually transmitted parasites are then able to invade the female’s organs to be further transmitted remains to be examined. Few studies have described sexual transmission of pathogens by soft or hard tick species. In experimentally infected *Ixodes persulcatus* and *Hyalomma anatolicum*, viral particles of the tick-borne encephalitis virus were detected by electron microscopy in spermatocytes and spermatids and the transmission of the virus from an infected male to a female was demonstrated, with its subsequent detection in the eggs (Chunikhin et al., 1983). The relapsing fever agent *Borrelia crocidurae* is also efficiently transmitted to uninfected females of *Ornithodoros erraticus* during copulation with infected males, with 23 and 37% of the females found infected respectively after the first and second gonotrophic cycle (Gaber et al., 1982). The specificity of this transmission was also demonstrated as *B. crocidurae* was not able to infect the testes of *O. savignyi* following an infective blood meal, while testes of *O. erraticus* fed on the same host were infected by the spirochaete (Gaber et al., 1984). The acquisition of *B. garinii* by *Ixodes persulcatus* females from naturally infected males is also suspected (Alekseev et al., 1999). The sexual transmission of *Babesia* sp. YLG could represent an important biological characteristic of clade V, but also an epidemiologically significant feature in the transmission cycle of this parasite. Due to the short seabird breeding season, acquisition/transmission cycles of the parasite need to occur within a few weeks, when adults as well as chicks are present in the nest. As *O. maritimus* females realize several successive gonotrophic cycles after copulation, sexually transmitted parasites may have time to spread within the female tick and invade salivary glands to be transmitted to the host, or ovaries to be transovarially transmitted. Increasing the ratio of infective females through sexual transmission could therefore be an efficient transmission strategy in soft ticks (several blood meals per life stage) compared to hard ticks (one blood meal per life stage).

In our previous study, we demonstrated the occurrence of *Babesia* sp. YLG two years in a row in chicks of the same breeding colony (Bonsergent et al., 2022) and, here, we find the parasite in ticks collected in this same location over four years. *Babesia* sp. YLG can therefore be considered as endemic to Carteau islet. The parasite reservoir may consist in *Babesia* infected ticks, surviving in the soil around the nests during the non-breeding period, or in infected adult gulls, or both. The prevalence of *Babesia* infected seabirds and parasite loads in the blood are usually lower in adults compared to chicks (Quillfeldt et al., 2014; Espinaze et al., 2019; Snyman et al., 2020), so the role of adult birds as reservoirs may be limited. However, further analyses of adult gulls are required, as many studies only use blood smears to detect blood parasites, a method that lacks the sensitivity required to detect asymptomatic carriers (Malandrin et al., 2004; Chauvin et al., 2009). Furthermore, parasitemia may change over time in infected birds, and particularly so during the pre-breeding period when a lot of energy may be allocated to reproduction and less to maintenance needs such as immunity (Gylfe et al., 2000).

*O. maritimus* may be a particularly good reservoir for parasites such as *Babesia* sp. YLG. Indeed, the life cycle of these ticks is relatively long – likely covering at least 2 breeding seasons, and individuals can remain in dormancy for many years when the host is absent (KD. McCoy, unpublished data). If transstadial transmission is efficient, this would allow ticks to locally maintain the parasite over long periods of time. The maintenance ability would be even higher if transovarial and sexual transmission pathways are verified in the system, up to the point where the parasite could be maintained without a required passage by the seabird host. However, regardless of the reservoir species, our previous study demonstrated a high transmission efficiency of *Babesia* sp. YLG to chicks, with 58-85% of infected during the first 3 weeks post-hatching, many with extremely high parasitemia (27 to 41% of individuals with parasitemia over 10%) (Bonsergent et al., 2022). Future work will now need to focus on the degree to which the parasite can be maintained without the vertebrate host and the epidemiological consequences of infection for both ticks and seabirds.

## Acknowledgements

We thank all the people that assisted with field sampling (Thomas Blanchon, Elodie Conte, Maxime Duhayon, Thibault Langlois, Louisianne Burkhart, Charly Souc, Florence Nono-Almeida) and provided excellent technical assistance in the lab (Caroline Hervet).

## Data, scripts, code, and supplementary information availability

Data are available online: 10.57745/XSSMO1 of the webpage hosting the data https://doi.org/10.57745/XSSMO1.

## Authors’ contributions

**Claire Bonsergent**: Investigation, Formal analysis, Methodology, Visualization, writing-Original draft preparation, Reviewing and Editing.

**Marion Vittecoq:** Resources, Funding acquisition, Reviewing and Editing.

**Carole Leray**: Resources, Reviewing and Editing.

**Maggy Jouglin**: Methodology, Reviewing and Editing.

**Karen McCoy**: Resources, Conceptualization, Funding acquisition, Writing, Reviewing and Editing.

**Laurence Malandrin**: Conceptualization, Funding acquisition, Supervision, Writing-Original draft preparation, Reviewing and Editing.

## Supplementary materials

### SM1

Location of 18S primers for the specific amplification of a 18S rDNA gene fragment of *Larus michahellis* (18SLm_fw and 18SLm_rev, green colored) and *Ornithodoros maritimus* (18SOm_fw and 18SOm_rev, orange colored) designed according to a Clustal Omega alignment. GenBank accession numbers correspond to the 18S rDNA gene sequence of *Babesia* sp. YLG (MZ541058), *Ornithodoros maritimus* (OP542591) and *Larus michahellis* (OP542589).

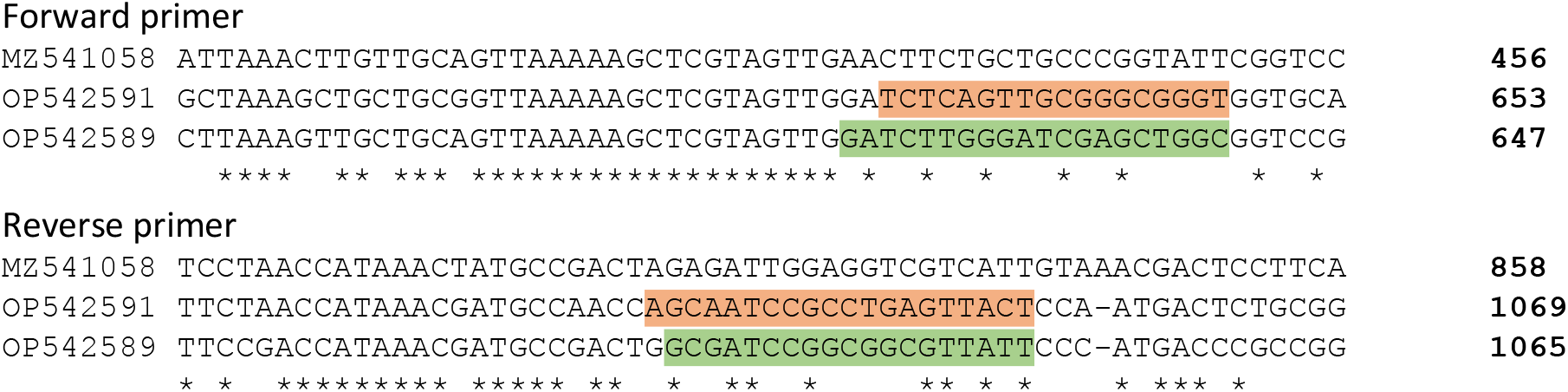

### SM2

Number of each organ type of *Ornithodoros maritimus* collected after dissection and used to test for the presence of *Babesia* sp. YLG (number of ticks in brackets).

| Sex/stage | Salivary glands | Ovaries | Endospermatophores | Male genitalia | Caeca |
| --- | --- | --- | --- | --- | --- |
| Females (107) | 103 | 105 | 66 | / | 101 |
| Males (15) | 12 | / | / | 11 | 14 |
| Nymphs (22) | 22 | / | / | / | 22 |

